# The role of network connectivity on epileptiform activity

**DOI:** 10.1101/2021.02.16.431388

**Authors:** Giuseppe Giacopelli, Domenico Tegolo, Michele Migliore

## Abstract

A number of potentially important mechanisms have been identified as key players to generate epileptiform activity, such as genetic mutations, activity-dependent alteration of synaptic functions, and functional network reorganization at the macroscopic level. Here we study how network connectivity at cellular level can affect the onset of epileptiform activity, using computational model networks with different wiring properties. The model suggests that networks connected as in real brain circuits are more resistant to generate seizure-like activity. The results suggest new experimentally testable predictions on the cellular network connectivity in epileptic individuals, and highlight the importance of using the appropriate network connectivity to investigate epileptiform activity with computational models.

## 1 Introduction

Epilepsy is a relatively common and widespread brain disease affecting people of all ages [1]. According to the World Health Organization, nearly 50 million people are affected by this disease [2]. For this reason, there are extensive experimental and theoretical efforts attempting to figure out the mechanisms underlying the onset and propagation of the plethora of abnormal (and transient) brain electrical activity caused by this disease. A better understanding of the involved mechanisms can facilitate the development of therapeutic solutions. A number of potentially important mechanisms have been identified as key players, such as genetic variations [3] often associated with ion channels mutations [4], alterations of synaptic function [5], and network connectivity [6,7]. However, current technical problems make the study of their specific contribution very difficult to investigate experimentally. From this point of view, computational models [8,9] can be a very convenient approach to identify the relative role and importance in generating seizures and, more generally, epileptiform activity.

Here we study in more details the role of network connectivity at cellular level, using a network composed of individual neurons implemented following a computational model, the Epileptor [10], which has been demonstrated to be able to reproduce epileptiform activity at the macroscale level, and it has been successfully used to simulate the possible scenarios after surgery in patients with drug-resistant seizures [11]. In this work, by exploiting a new mathematical framework [12,13] that can create networks with connectivity properties similar to those observed in real brain networks, we studied how network connectivity can affect the onset of epileptiform activity. The model suggests that networks connected as in real brain circuits are more resistant to generate seizure-like activity. The results suggest new experimentally testable predictions and highlight the importance of using the appropriate network connectivity to investigate epileptiform activity with computational models.

## 2 Methods

### 2.1 Single neuron model

The first step in building a network is to decide on the type of individual neurons composing it. We decided to start from the Epileptor [10] (since now EP), which has been shown to be able to reproduce epileptiform activity.

The original model uses six differential equations to represent the activity of a population of Hindmarsh–Rose (HR) neurons [14], under a mean-field approximation ([15], defined as the average of the membrane potential associated to neurons). In order to more easily implement different connectivity rules, we modified this model by using independent equations for the excitatory and inhibitory neurons, and by adding units to the original adimensional implementation. The membrane potential of an excitatory neuron, *x*_*Ei*_, was thus defined as:

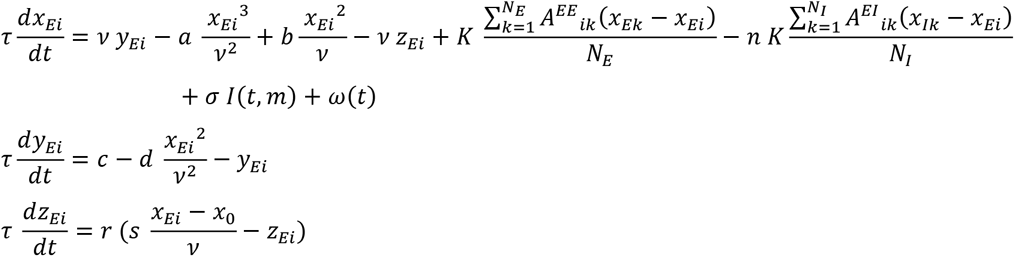

Analogously, the equations for an inhibitory neuron *j* were defined as:

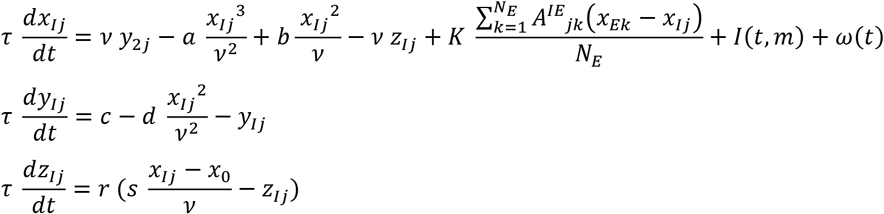

Where *x* is the membrane potential (in µV) of neuron *x, y* and *z* are the fast-slow variables of the original EP model, *N*_*E*_ and *N*_*I*_ are the number of excitatory and inhibitory neurons, respectively, the parameters *a=1, b=3, c=1, d=5, s=4, r = 0*.*003, x*_*0*_*=-1*.*6* and *K=150* define the fast-slow interaction, and n is the inhibitory strength. The matrices *A*_*EE*_, *A*_*EI*_ and *A*_*IE*_ are the adjacency matrices for each type of connection (Excitatory-Excitatory, Excitatory-Inhibitory and Inhibitory-Excitatory; Inhibitory-Inhibitory connections were ignored, as in [8]). The variables *τ=0*.*0167 s, ν=20 µV* and *σ=1 µV/nA* allowed us to introduce units to the model. The function *ω(t)* is a noise variable and *I(t,m)* is the time- and space-dependent external current.

### 2.2 Theoretical background for network connectivity

Neurons’ connectivity can be encapsulated in the concept of spatial graphs [13]. A spatial graph *G* is an abstract entity composed by a set of vertices *V* (in our case, the neuron somas), a set of edges *E⊆ V*×*V* (in our case, the synaptic connections), and a set of positions (in our case, the spatial coordinates of the somas). Each vertex is associated with indegrees (the number of incoming connections) and outdegrees (the number of the outgoing connections). The set of degrees is described by the probability density function of their distribution. Finally, a connection length distribution can also be associated to a network with spatially distributed nodes.

Neuronal networks are very often connected following either a trivial all-to-all rule or according to fixed probabilistic rules based on two main graph-theoretical models: power law or exponential. Power law models have a heavy tail behavior in the degree distributions [16], whereas exponential models [17] (since now ER) use a fixed connection probability rule that generates an exponential decay of the degree distributions around a central value. Real brain networks are not connected all-to-all, but exhibit a mix of exponential and power law connectivity models: an exponential behavior for low degrees, and a power law behavior for high degrees.

We have recently mathematically demonstrated how an appropriate convolution of these models can quantitatively reproduce the degree distributions observed in a real brain network [12,13]. The key idea of the algorithm behind the theory is to split a network in blocks, and use a heavy tail connectivity inside the blocks and an exponential connectivity between blocks. In detail, assuming that Ω_A_ and Ω_B_ are two spatial regions with *N*_*A*_ and *N*_*B*_ neurons, respectively, within each region a power law tail connectivity is created using a growing network algorithm [18] with a cost function defined as

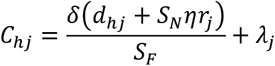

where *δ* and *η* are parameters of the model, *d*_*hj*_ is the square of the Euclidean distance between nodes *h* and *j, λ*_*j*_ is a graph theoretical measure of centrality [18], *r*_*j*_ is a uniform random number in [0,1], *S*_*N*_*=200 µm*^*2*^ and *S*_*F*_*=1 µm*^*2*^. If *f*^*I*^_*k*_ and *f*^*O*^_*k*_ are the limit distributions of this process, assuming an ER connectivity between blocks, and defining the graph *C* as the union between *A* and *B*, it can be mathematically proved [12,13] that

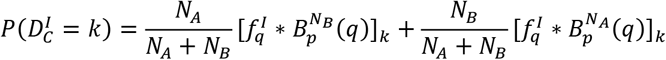

Where *P(D^I^_C_=k)* is the indegree distribution of graph*C, B_p_^N^=(k)* is the binomial distribution and *\** is the convolution operator defined between two discrete distributions *h*_*k*_ and *g*_*k*_ as

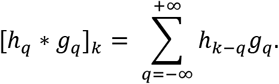

The outdegree distribution is described in the same way. A web application allowing to create an arbitrary network with the desired distributions will be available in the live papers section of the Human Brain Project (HBP) (https://humanbrainproject.github.io/hbp-bsp-live-papers/index.html), and the python code with model and simulation files will be downloadable from ModelDB (https://senselab.med.yale.edu/modeldb/, acc.n. 266910).

### 2.3 Simulations

For all simulations we used networks of 550 neurons (500 excitatory and 50 inhibitory) connected using four different topologies: an exponential connectivity rule, with or without a distance-dependent connection (ER and ER-dist models), or a convolutive model reproducing the in- and outdegree distributions experimentally observed in a *C. elegans* brain [19] or in a mouse hippocampus slice [20]. For simulations where the connection length distribution was also considered, neurons were randomly distributed in a 3D volume shaped either as a rectangular *500×500×2000 μm* volume (for *C. elegans* connectivity), or as a *400×300×10 μm* slice (for hippocampal connectivity). All simulations were implemented using Matlab [21], and model files will be available on ModelDB (acc. n.266910). Following a custom practice for this type of electroencephalographic (EEG) experimental recordings, simulation traces were filtered with high-pass (1 Hz) and a low-pass (40 Hz) Butterworth filters [22]. This allowed a better comparison between modeling and experimental results.

To activate the networks, inputs were implemented as external currents with a peak value located at the center of the model, and modulated according to the spatial soma locations as:

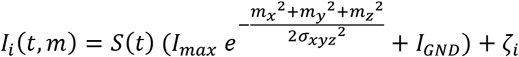

Where *{mx, my, mz}* are the i-th soma coordinates, *I*_*max*_ *= {1*.*5nA, 2*.*5nA, 5nA}, I*_*GND*_*=1nA, S(t)* is the time-dependent component of the current, with values between 0 and 1, *σ*_*xyz*_*=600 µm*, and ζ_i_ a random normal vector with amplitude 0.1 nA.

In Figure 1a we report a typical normal EEG recording (from [23]), and in Fig. 1b a trace exhibiting epileptiform activity from a public database [24], together with the corresponding Welch spectrograms [22]. The epileptic trace shows the characteristic high-amplitude population spikes and a significant increase in the spectral density in the low frequency range (compare right plots of Fig. 1a and 1b). A typical model trace, obtained from the latest Epileptor version [11], and the corresponding spectral density is shown in Fig. 1c. It demonstrates that this type of model is able to reproduce epileptiform activity and spectral properties in good qualitative agreement with experiments. In order to have a quantitative measure of an epileptiform activity, we used the average network activity, calculated from the average potential of the neurons [10] as:

**Fig. 1.**
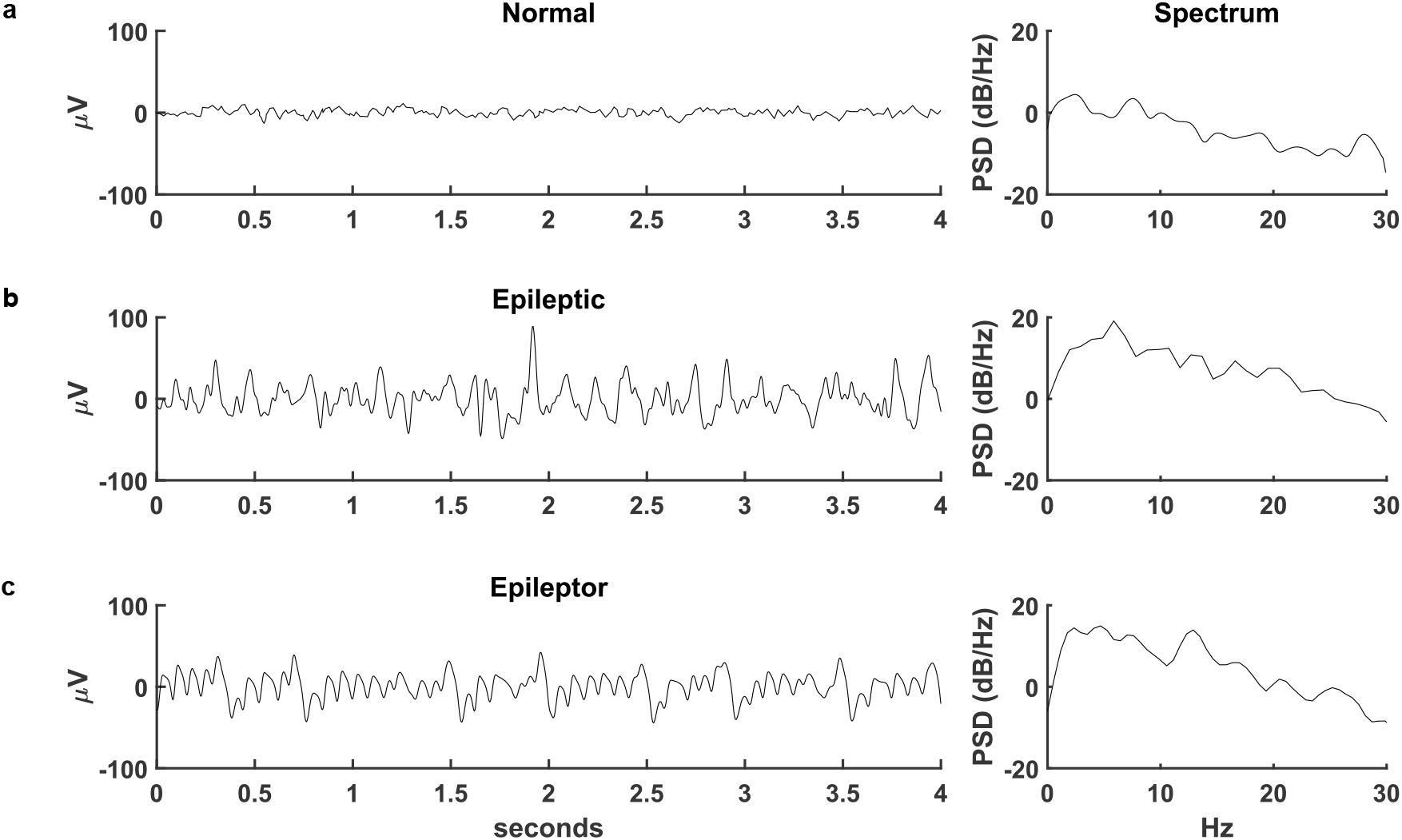
Comparison of experimental and model traces. A) typical EEG trace under control conditions (Left, from Fattinger 2017), and the corresponding Welch spectrum (Right); B) typical EEG trace during epileptiform activity (from Zwoliński 2010); C) time course of the variable corresponding to the membrane voltage during a simulation of an Epileptor (El Houssaini 2020) with parameters: *a = 1, b = 3, c = 1, d = 5, I*_*ext1*_ *= 3*.*1*, m = 0, *a*_*2*_ *= 6, τ*_*2*_ *= 10, I*_*ext2*_ *= 0*.*4, γ = 0*.*01* and a white noise of amplitude 0.25. The adimensional model has been converted in a dimensional one using the time scale constant 0.0167 s and the voltage scale constant 25 µV.

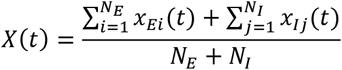

In the example traces shown in Fig. 1, the average potential was 6.9, 29.9, and 30.8 µV for the normal, epileptic, and Epileptor trace, respectively. For the rest of this paper, we will define a trace as exhibiting an epileptiform activity when the average membrane potential is above 18.4µV, half-way between the values calculated from the normal and epileptic experimental traces.

## 3 Results

In order to investigate the role of network connectivity in the onset of epileptiform activity, we considered two aspects: 1) the type of connectivity (exponential or convolutive), 2) the spatial arrangement of the individual neurons in a volume, shaped as a column (like in the cortex) or as a slice (like in the in vitro hippocampal preparation). The rationale for this choice was that we were interested in studying the response of networks connected using a convolutive model, and compare the results obtained with networks implemented using the widely used ER model, in which neurons are connected with a fixed connection probability. We think that this is important, because it has been demonstrated [12,13] that a convolutive model better represents the connectivity of a real brain network, and can thus provide a better insight into the network mechanisms responsible for the onset of epileptiform activity in the real brain.

In Fig. 2a we plot the degree and connection length distributions of the networks implemented by uniformly distributing the 550 Epileptors in a rectangular volume, connected in three different ways: 1) following an ER model, ignoring the distance among neurons (Fig. 2a, red traces), and with a fixed connection probability of *p=0*.*026*, corresponding to the average connection probability of a *C. elegans* brain, 2) following an ER model but with a connection probability depending on the distance between neurons as *p(d)=Ae*^*-Bd*^ (as in [25]), where *d* is the distance between two nodes, *A=0*.*2*, and *B=0*.*004* (Fig. 2a blue traces), and 3) using a convolutive model (Fig. 2, green traces) fitting a *C. elegans* degree distributions [19], with two blocks and parameters, *δ=1*.*5, Ek=1* and *η=3* (see Methods). The degree distributions using the ER models (Fig. 2a red and blue traces) were very similar to each other, even if their connection length distributions were very different (compare red and blue lines in the right panel of Fig. 2a). This suggests that modifying the connection probability by including information on the distance between neurons does not significantly change the degree distributions. The qualitative difference, between the ER models and the convolutive model, in the connectivity among neurons, can be better appreciated using a graphical representation as in Fig. 2b, where we present a circular graph representing the connectivity of 50 randomly chosen individual neurons from each network. Note the relatively dense and uniform connectivity of neurons belonging to ER-type networks (Fig. 2b, left and middle plots), with only very few neurons poorly connected with the other neurons in the plot. This is in striking contrast with the sparser and distributed connectivity exhibited by the convolutive model networks, with a large proportion of poorly connected neurons and a significantly higher proportion of hub neurons.

**Fig. 2.**
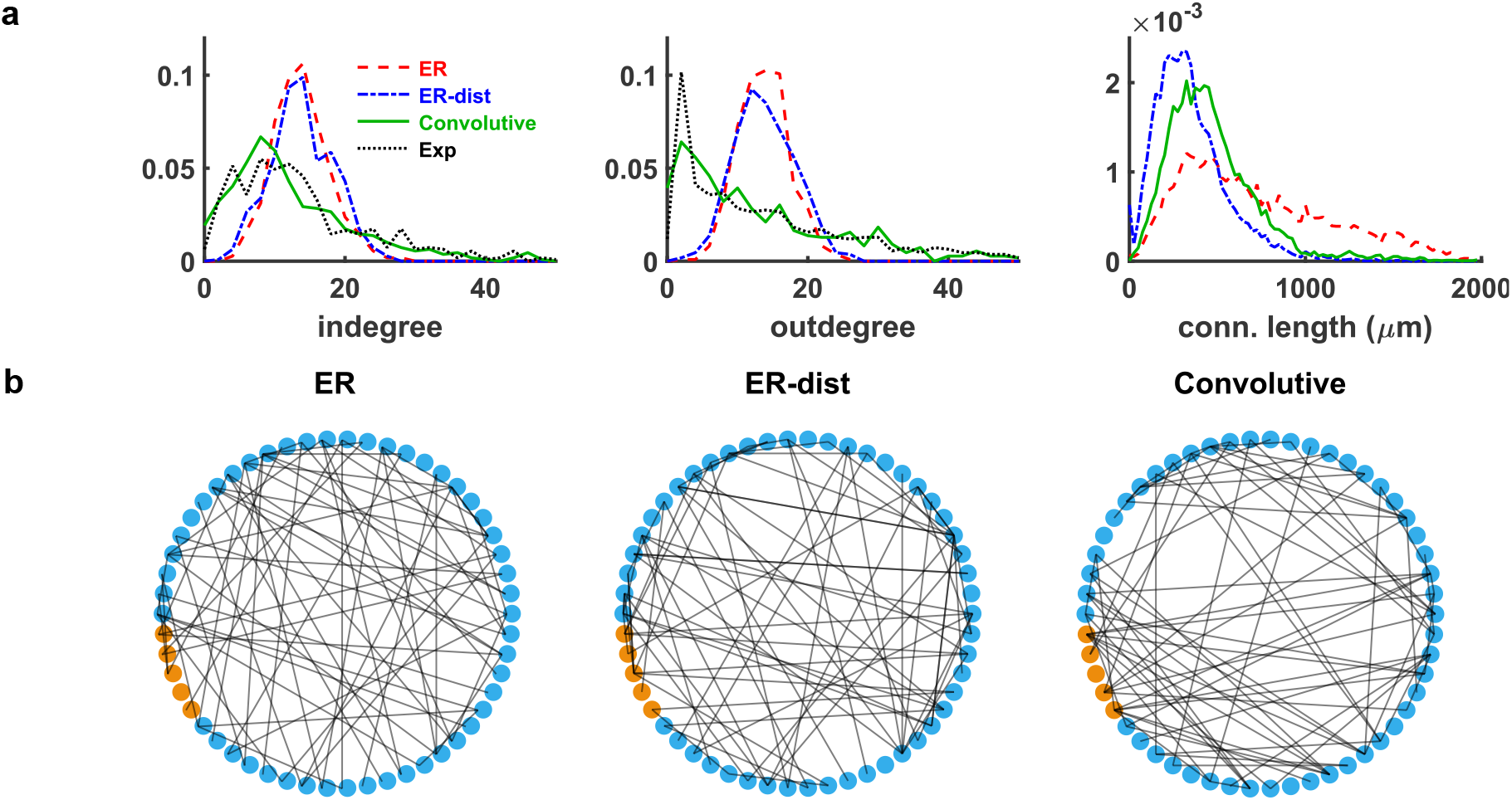
Neurons distributed in a thick rectangular volume. A) Indegree distribution (left), outdegree distribution (middle) and connection length distribution (right) for networks of Epileptors connected as an ER model (red), ER with distance (blue) and using a convolutive model (green) fitted to experimental *C. elegans* data (black, Ref. 19, connection length was not available). B) Circular graph representation of connectivity for 50 randomly chosen neurons for the ER model (left), the ER-dist model (middle) and Convolutive model (right). Blue and orange circles represent excitatory and inhibitory neurons, respectively.

The results did not qualitatively change when we considered networks of neurons distributed in a volume shaped as a slice, i.e. a rectangular volume of *400×300×10 μm*, with an average connection probability (*p=0*.*0438*) consistent with that observed in a hippocampal slice, as shown in Fig. 3. The ER model (Fig. 3a, red), and the ER-dist model (Fig. 3a, blue) with parameters *A=0*.*7* and *B=0*.*025*, still had a relatively similar degree distribution, in spite of a very different connection length distribution (Fig. 3a, right). Both were rather different from the results obtained with the convolutive model (Fig. 3a, green) by fitting (*p>0*.*05*) the experimentally observed degree distributions (Fig. 3a, black). The connection length distribution reflected the very different shape and size of the volume in this case (note the different scale for the distance axis in Fig. 2a and Fig. 3a), with a proportion of longer connections for the convolutive model more similar to the ER model built without distance-dependent connectivity. The circular graph representation in Fig. 3b, reinforces the contrast between the dense and uniform connectivity of neurons belonging to ER-type networks and the clear predominance of hub neurons for the convolutive model. This result is consistent with experimental findings on functional network connectivity at macro scale during seizures [6]. Taken together, these results highlight the large overall structural differences between networks with a realistic connectivity and those built using a fixed connection probability, independently from the network shape and size. This is a key difference and, as we will see in the following paragraphs, it suggests that a network connected as real brain network can be significantly more resistant to generate an epileptiform activity, with respect to networks with a fixed connection probability.

**Fig. 3.**
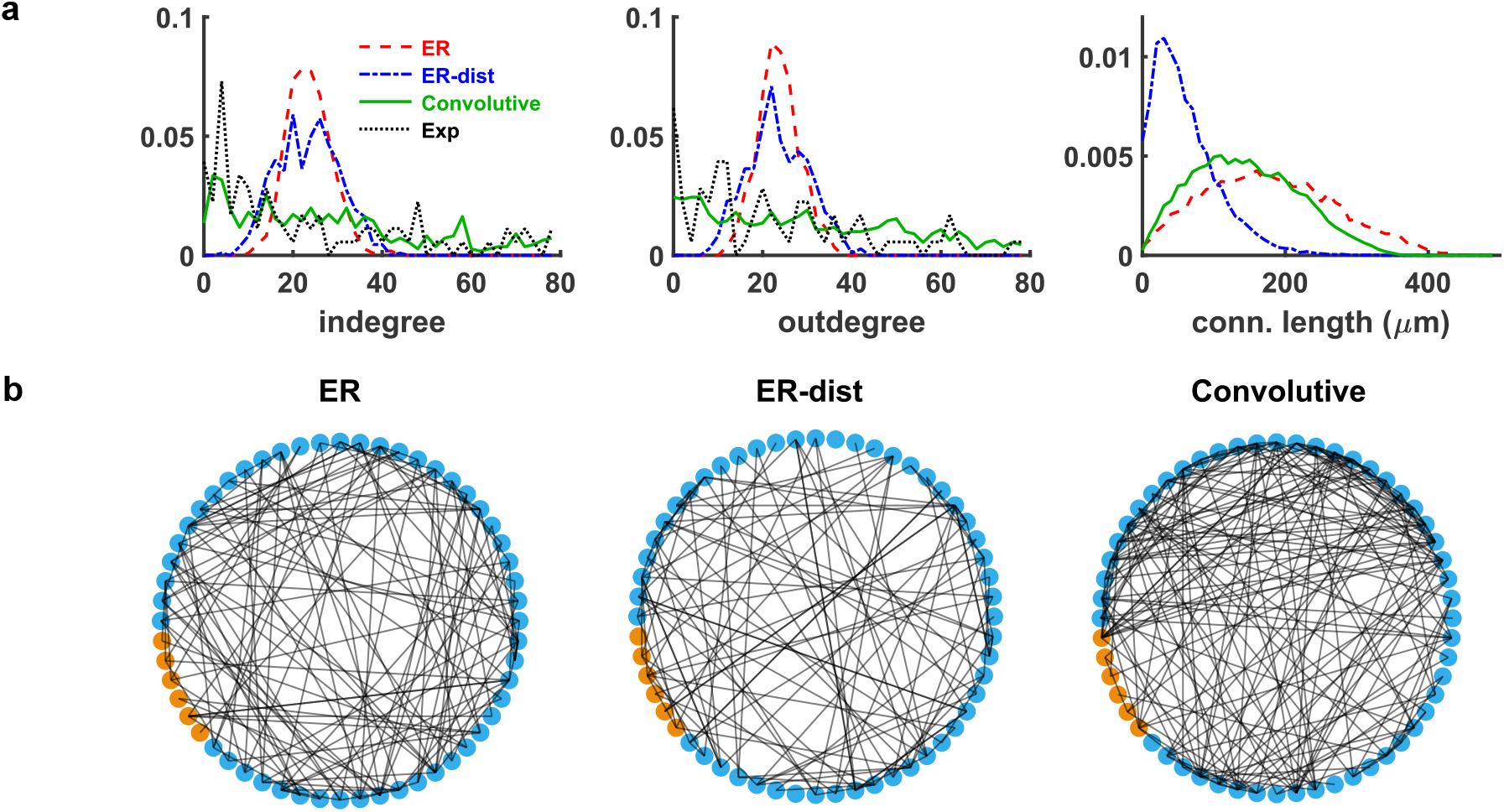
Neurons distributed in a thin rectangular volume. A) Indegree distribution (left), outdegree distribution (middle) and connection length distribution (right) for networks of Epileptors connected as an ER model (red), ER with distance (blue), and using a convolutive model (green), connected as observed in an experimental hippocampal slice (black, Bonifazi 2009), implemented with 4 blocks and 2 sub-blocks containing 1% and 99% of the neurons, respectively, and with parameters *δ=0*.*25, Ek=0*.*1* and *η=3*. B) Circular graph representation of connectivity for 50 randomly chosen neurons for the ER model (left), the ER-dist model (middle) and Convolutive model (right). Blue and orange circles represent excitatory and inhibitory neurons, respectively.

### 3.1 Networks with realistic connectivity are less prone to generate epileptiform activity

To test the response of our networks to an external input, we carried out a series of 16 sec simulations during which we activated, on top of the background activity, an external current, spatially modulated as described in the Simulations section, with *I*_*max*_ *=5nA* and the time course shown in Fig. 4a. In Fig. 4b and Fig. 4c we plot the response of the networks, in terms of average membrane potential over all neurons, under different conditions of spatial distributions and connectivity rules. All networks received the same input. Before the stimulus onset, networks (with or without distance dependent connectivity) started to have an epileptic behavior, with an average value in the 5-15 sec time window of 46.6, 34.6, 47.3 and 37.1 µV, for the ER and ER-dist in thick volume and for the ER and ER-dist in thin volume, respectively. Networks connected as in real brain networks (Conv. Model in Fig. 4b-c), had an average value of 9.9 and 7 µV, for thick and thin volumes, respectively, very close to the value obtained from normal traces. These results suggest that the structural connectivity of real brain networks is more resistant to enter into an epileptic state.

**Fig. 4.**
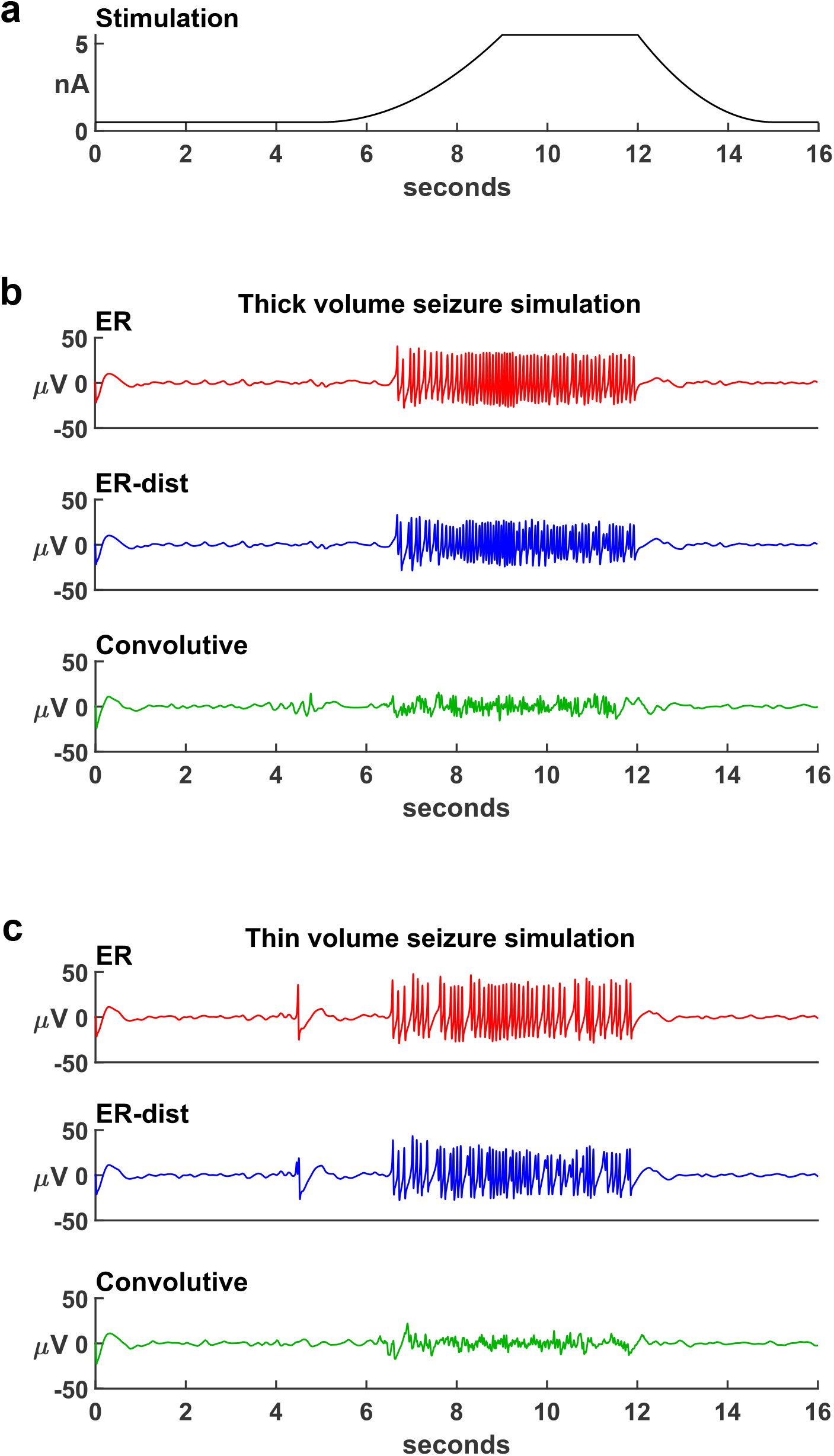
Networks with realistic connectivity are less prone to generate epileptiform activity. A) Maximum input current, in the center of the nework, as a function of time; B) Simulation of epileptiform activity in networks of Epileptors distributed in a thick rectangular volume and connected in different ways, with low inhibitory interaction (*n=0*.*1*). C) Same as in B) but with neurons distributed into a thin rectangular volume. the activity in all cases appeared to have a normal behavior, with occasional and short-lived spikes of higher intensity, more noticeable for the ER models in thin volume. As the external input increased, all ER model

### 3.2 Realistic connectivity robustly protects from the consequences of strong inputs

In addition to the input strength, the amount of inhibition generated by network activity is a well-known modulatory mechanism for epileptiform activity. Thus, we explored this aspect by carrying out a series of simulations in which we progressively increased the parameter *n*, regulating the inhibitory interaction among neurons. Simulations were performed for three values for *I*_*max*_ and a discrete set of *n* values (0.05, 0.1, 0.15, 0.2, 0.3, 0.4, 0.5, 0.75, 1.0 and 1.5). For each combination, we simulated five network instances. In Fig. 5, we show the mean amplitude during the stimulation for each model. As expected, increasing the inhibition will eventually reduce the average amplitude of the signal. However, large values are not physiological. If we focus on the lower range of *n* values (e.g. below 0.5) it can be observed that simulations using the Convolutive model have a mean amplitude lower than exponential models at all currents (Fig. 5, top panels). It should also be noted that, for high input currents, ER distance models have lower mean amplitudes than ER models, pointing out the importance of spatial information in building networks models. The same trend can be observed in the bottom panels of Fig. 5, for a thin volume model. These results suggest that a realistic connectivity, implemented though a convolutive model, robustly protects from the onset of epileptiform activity during a wide range of strong external inputs.

**Fig. 5.**
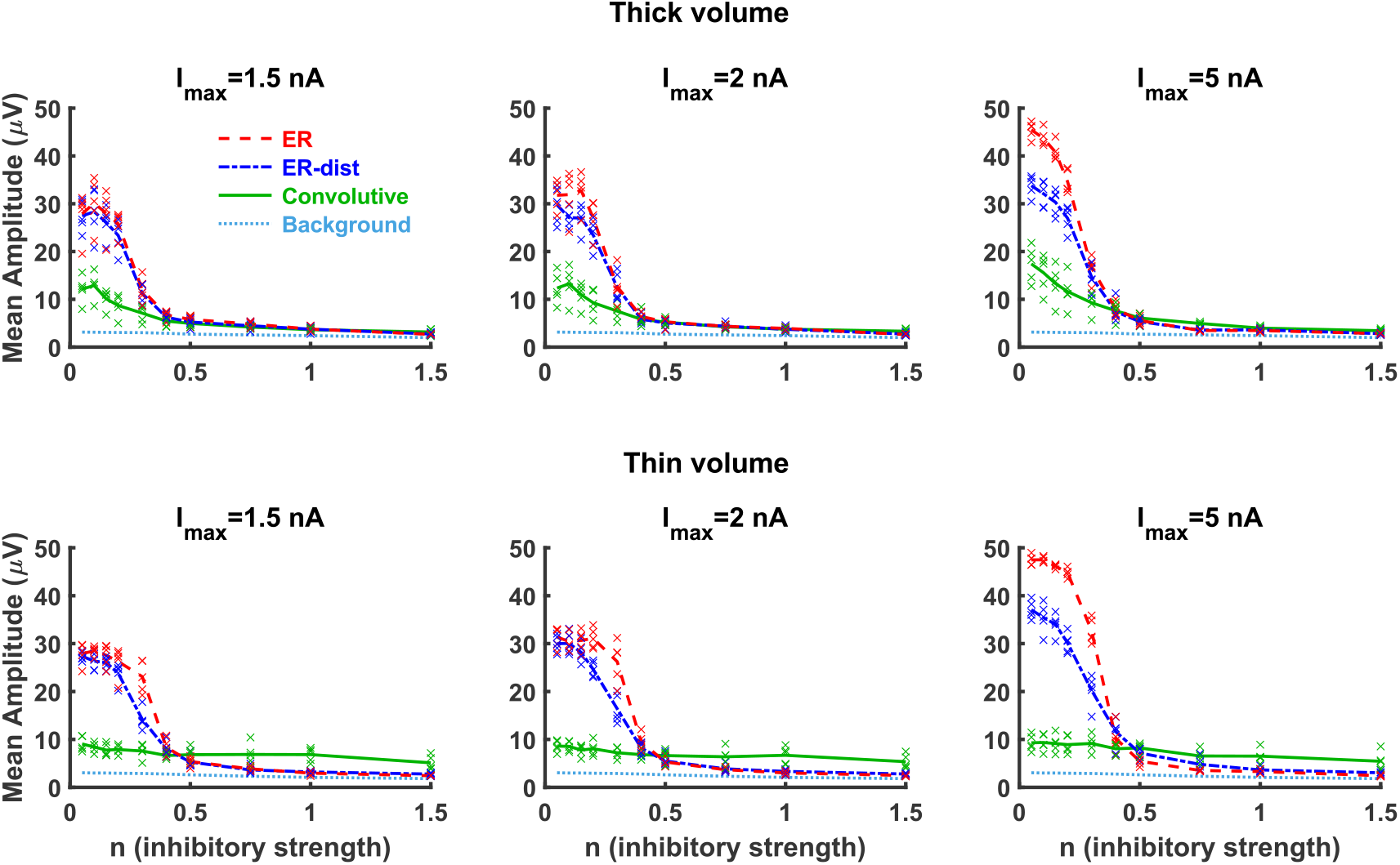
Realistic connectivity robustly protects from the consequences of strong inputs. Average membrane potential of the different models calculated, as a function of the inhibition, over the 10 sec duration of an external input of different maximum strengths, *I*_*max*_. For any given value of *I*_*max*_ and *n*, symbols represent the average membrane potential obtained from five network instances. Lines represent their average value.

### 3.3 Spike-wave complexes

Epileptiform activity includes spike-wave complexes [26], which are identified as EEG portions with variable amplitude and frequency [27]. A typical spike-wave complex period is shown in Fig. 6a. With our Epileptor networks, we found that they can be modeled by assuming a dynamic modulation of the inhibitory strength. This corresponds to the physiological plausible assumption that the inhibitory interaction among neurons follows network activity (in terms of average membrane potential), generating a dynamic level of inhibitory response. We implemented this effect by modulating the value of *n*, the variable responsible for the inhibitory interaction among Epileptors, as:

**Fig. 6.**
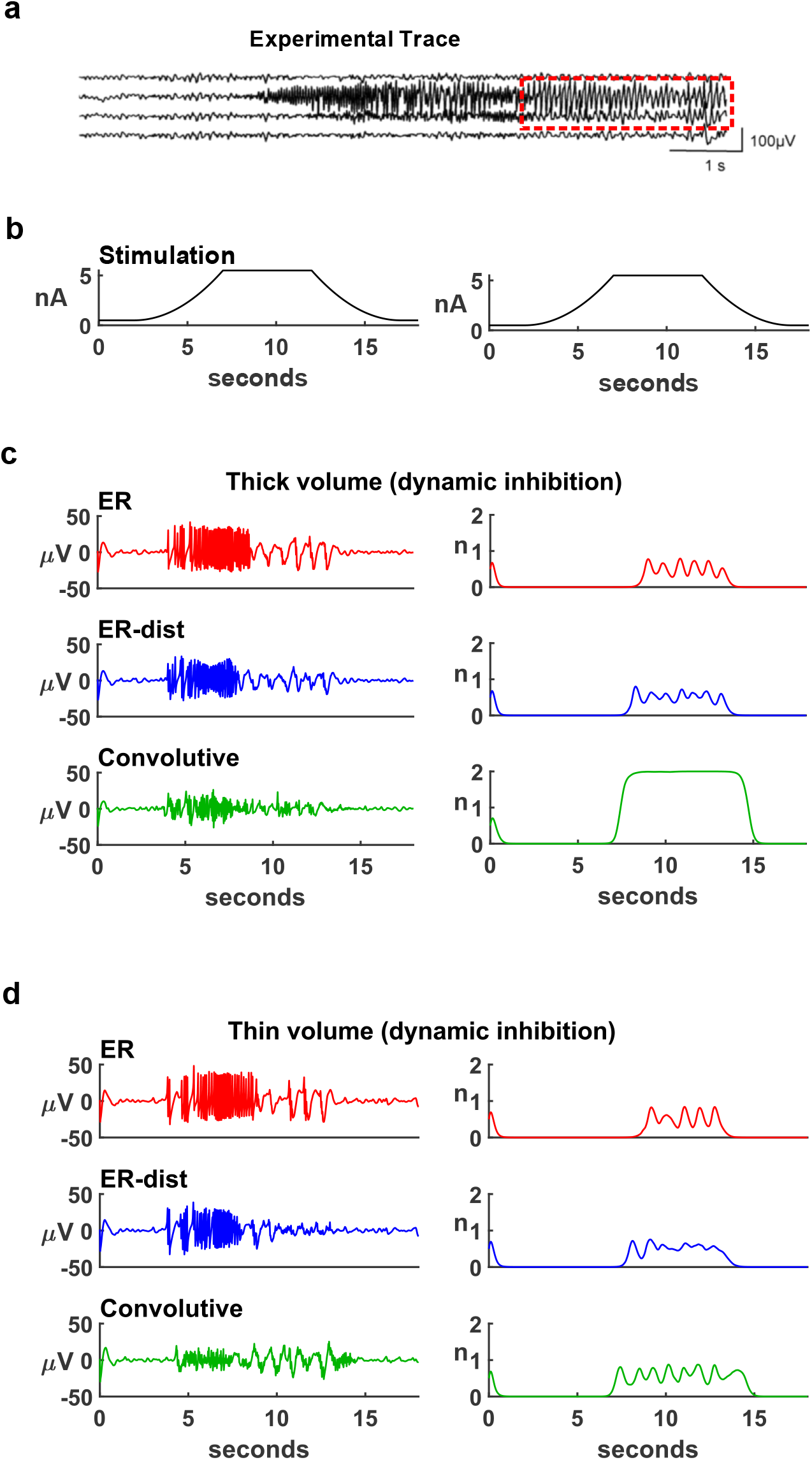
Dynamic inhibition can underlie spike-wave complexes. A) Experimental EEG recording showing a period of spike-wave complexes (from Noachtar and Rémi 2009), indicated by the red outline; B) Maximum input current, in the center of the network, as a function of time; C) Average membrane potential for neurons distributed in a thick volume (left plots) and time course of n, the variable associated with the inhibitory interaction (right plots), for: ER model (top row), ER distance model (middle row) and Convolutive model (bottom row). D) Same as in C) but for neurons distributed in a thin volume.

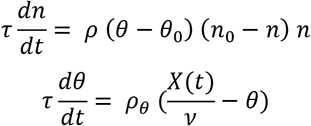

where *τ=0*.*0167 s, ν=20 µV, n*_*0*_*=2, ρ=0*.*05, ρ*_*θ*_*=0*.*15* and *θ*_*0*_ *= {-0*.*9; −1}* for the thick or the thin volume, respectively. For these simulations, the input stimulus time course is shown in Fig. 6b, and the time course of the average membrane potential for the different models and neurons distributed in a thick volume in Fig. 6c (left plots). Spike-wave complexes are clearly evident in the ER models (Fig. 6c, red and blue traces), following a period of a seizure-like pattern. The average values during the stimulation period (3-17 sec), were 29.8 and 20.5 µV. Spike-wave complexes were also present for the convolutive model (Fig. 6c, green trace) but they were of much smaller amplitude, and the overall average activity during the stimulation period was within the range for a normal activity (11.8 µV).

Interestingly, the inhibitory interaction for the convolutive model was much stronger than for the ER models (Fig. 6c, right plot), indicating that a convolutive model connectivity is able to maintain network activity within normal levels better than ER networks. This latter aspect explains why in networks of neurons distributed in a thin volume (Fig. 6d), where the overall connectivity is reduced, spike-wave complexes are evident also for the convolutive model (Fig. 6d, green traces), although the average network activity during the stimulation (8.1 µV) is still within a normal level, and much smaller than for ER and ER-dist models (31.7 and 19.2 µV, respectively). Taken together, these results suggest that spike-wave complexes can emerge from a dynamic inhibitory interaction among neurons.

## 4 Discussion

The results presented in this paper point to a paramount role for network connectivity in modulating the onset of epileptiform activity during strong localized inputs, which can arise during a variety of physiological brain states or cognitive activities [28]. The relevance of these findings can be discussed in the context of both modeling and experiments.

From the modeling point of view, it should be stressed that large-scale neuronal network models, aiming at studying the onset and propagation of epileptic activity, so far have been almost exclusively implemented using all-to-all or fixed connection probabilities. However, these connectivity rules do not represent the real connectivity experimentally observed in brain networks. We have previously shown [29] how this difference can generate significant variations in the response of a network to a given input. Here, using a recently published mathematical framework, able to create convolutive model networks reproducing experimental findings [12,13], we have shown that networks connected in the same way as real systems are less prone to generate epileptiform activity. Furthermore, the model’s suggestion that spike-wave complexes can be generated by a dynamic inhibitory interaction, also points to a possible improvement on how to reproduce this specific experimental feature in a model. These results thus suggest that network models should be built using physiologically plausible connectivity rules, rather than fixed connection probabilities. This will make it easier to interpret experimental findings in terms of model parameters and it will allow making more precise and successful, experimentally testable, predictions.

From the experimental point of view, our results suggest that epileptiform activity can arise from an epileptogenic process during which a network (or a significant part of it) is progressively transformed from normal to epileptic through an ictogenetic mechanism that modifies the connectivity distribution functions (i.e. in/outdegrees and connection length) from a convolutive to an exponential scheme. This process is equivalent to selectively delete hub neurons from the network, and it makes the network less robust and more likely to generate epileptiform activity in the presence of strong inputs. There are a number of experimental findings indirectly supporting this result (reviewed in [1]), which may be the consequence of pathogenetic mechanisms altering synaptic plasticity and axonal sprouting.

Finally, the model suggests two specific experimentally testable predictions. The first one is that epileptic tissue should have a distinctly different cellular connectivity, with respect to normal tissue: there should be less hub neurons in the epileptic tissue. The second one is that spike-wave complexes can be modulated by a dynamic inhibition much higher than during normal activity. The role of inhibition in controlling epileptic activity is well known, and its generalized increase is the classic strategy followed by antiepileptic drugs [30]. However, this action has a wide range of negative collateral effects [31]. The model suggests that better results could be obtained by selectively increasing inhibition only when and where needed, i.e. in highly active regions. Under this condition, synapses are presumably activated at a higher rate. Pharmacological applications, acting above a threshold activation level on the biochemical processes underlying short-term facilitation of inhibitory synapses, could increase the amount of signal summation only during high-frequency synaptic activation in localized regions, generating less collateral effects in the rest of the brain. From this point of view, there are already experimental indications that specific proteins or compounds can regulate synaptic transmission, such as synapsins, which can regulate GABA release [32], and antiepileptic drugs can alter short-term plasticity [33].

## Acknowledgments

This research has received funding from the European Union’s Horizon 2020 Framework Programme for Research and Innovation under the Specific Grant Agreement numbers 945539 (Human Brain Project SGA3). Editorial support was provided by Annemieke Michels of the Human Brain Project.

